# GBA Solver: a user-friendly web server for growth balance analysis

**DOI:** 10.1101/2025.05.14.653971

**Authors:** Sajjad Ghaffarinasab, Hugo Dourado

**Affiliations:** Institute for Computer Science and Department of Biology, Heinrich Heine University, Universitätsstraße 1, D-40225 Germany

## Abstract

GBA Solver is a web-based application that enables efficient construction and numerical solution of coarse-grained growth balance analysis (GBA) models containing up to 20 reactions. The platform features an intuitive spreadsheet-based interface that eliminates the need for programming, integrates kinetic parameters from the BRENDA enzyme database, and provides interactive visualizations for model exploration. By lowering technical barriers, GBA Solver makes GBA modeling accessible to a wider scientific community, expanding the computational toolkit for investigating cell metabolism and growth. GBA Solver is freely available at: https://cellgrowthsim.com/. The source code is available at: https://gitlab.cs.uni-duesseldorf.de/general/ccb/GBASolver.

## 1. INTRODUCTION

Computational models are essential for studying the complex biochemical processes that govern cell growth and metabolism. Growth Balance Analysis (GBA) provides a simplified modeling framework for studying the optimal resource allocation in kinetic models that integrate metabolism, protein synthesis, and the dilution of all cell components at balanced growth [6, 7]. Different from other modeling frameworks, GBA does not require biomass composition as input; instead, it predicts the entire cellular composition by assuming optimal growth of a self-replicating system subject to mass conservation, nonlinear kinetic rate laws, and total density constraints. GBA outputs the cell growth rate, fluxes and biomass composition at different environments by solving a nonlinear optimization problem which requires as inputs a stoichiometric matrix, kinetic parameters and cell density. The numerical solution of such optimization problems are however computationally expensive and require programming skills. Here we present GBA Solver, a webbased application that enables efficient construction and numerical solution of coarse-grained GBA models. The platform provides an intuitive interface that eliminates the need for programming, making GBA more accessible to a wider community of researchers.

### A. Platforms for modeling cell metabolism and growth

There are currently several platforms and frameworks available for modeling cell metabolism, each with different levels of detail, mathematical formalisms, and usability. For linear, constraint-based workflows such as flux balance analysis (FBA) [22] and its extensions, tools like CNApy [28], Escher [15], ModelExplore [19], CAVE [18], and Fluxer [11], provide convenient model construction and visualization. For nonlinear kinetic models, software applications include COPASI [12], Tellurium [4], Virtual Cell [25], and AMICI [9], and web-based environments such as JWS Online [21] and runBioSimulations [27]. To contribute to the community and extend this ecosystem, we developed the GBA Solver, a web-based implementation of the GBA formalism for nonlinear self-replicating cell models. GBA models are unique in the sense they are nonlinear kinetic models that incorporate at the same time 1) the necessary production of proteins to catalyze each reaction in the model, 2) the dilution of all cell components by growth, including proteins and metabolites, and 3) a limited cell density including both metabolites and proteins. This approach allows GBA to predict the whole biomass composition (including metabolites and proteins) instead of requiring it as a modeling input, thus capturing global trade-offs in cell resource allocation without extra phenomenological assumptions. One downside of this approach is, as for any other type of kinetic model, the extreme computational expense to solve large nonlinear problems numerically. Here, we focus on a computational tool for GBA models including up to 20 reactions, which is sufficient to capture major cell functions in a coarse-grained manner [5, 10, 13, 20, 26, 29].

### B. Coarse-grained cell models

Coarse-grained models are useful tools for studying fundamental principles of cell physiology. By simplifying complex networks into a few effective reactions and catalytic sectors, these models can capture important trade-offs, such as the growth law of ribosome allocation [8, 26], shifts in metabolic strategies [20], and proteome partitioning [13], without requiring exhaustive mechanistic detail [5, 10, 13]. This type of simplification renders nonlinear optimization problems tractable while still providing insight into global resource allocation strategies [8, 20, 26, 29]. In this spirit, the GBA Solver is designed for coarse-grained self-replicator models, enabling the rapid, interpretable exploration of how metabolite concentrations and growth-driven dilution shape cell behavior.

## II. DESIGN AND IMPLEMENTATION

GBA Solver is built using the Shiny package [3] in R (version 4.4.1) [24]. The front-end architecture integrates HTML for content structure, CSS for visual styling, and JavaScript for dynamic interactions and enhanced client-side performance based on the Mazer dashboard template. We integrate comprehensive enzyme kinetic data from the BRENDA database [2], providing users access to curated turnover numbers *(k*_cat_) and Michaelis constants (*K*_m_) filtered to include only wild-type enzyme parameters, which are then systematically organized by Enzyme Commission (EC) numbers. Through an interactive table powered by the reactable package [17], users can efficiently search and filter enzyme parameters based on multiple criteria including organism, EC number classification, and substrate specificity. This streamlined interface simplifies the often challenging task of identifying appropriate kinetic parameters for metabolic models.

### A. Input data structure and model configuration

GBA models account for kinetic rate laws by relating the reaction flux *v*_*α*_ of each reaction *α* with the respective concentration *p*_*α*_ of the protein catalyzing that reaction (transport, enzymatic, or “ribosome” reaction producing proteins) and a turnover time function *τ*_*α*_ = *τ*_*α*_(**a, c**)

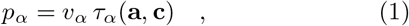

where **a** is a vector of external environment concentrations, and **c** is a vector of internal metabolite concentrations, both in units *gL*^*−*1^. The functions *τ*_*α*_ also depend on kinetic parameters, which is the simplest case of a Michaelis-Menten kinetics are the turnover number (*k*_cat_) and the Michaelis constants (*K*_m_). GBA Solver accounts for a larger set of possible kinetic rate laws by implementing the “convenience kinetics” [16], a general framework that can describe either irreversible or reversible rate laws under regulatory effects from activators and inhibitors. This includes, for example, an irreversible reaction *α* under some regulatory effects from activators, encoded by the function

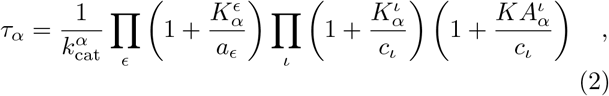

where the kinetic parameters are the (forward) turnover number 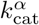 of that reaction *α*, the Michaelis constants 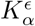 for external reactants *ϵ* involved on that reaction *α*, the Michaelis constants 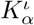 for internal reactants *ι* involved on that reaction *α*, and the activation constants 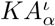 that quantify activator effects of internal reactants *ι* on that reaction *α*.

In GBA Solver we organize all the possible model parameters conveniently as matrices in different tabs of an open-source spreadsheet format (ODS), consisting of the following:

- Mass fraction matrix **M** (including external and internal reactants): quantifies the mass fraction of each reactant (row in the matrix) going through each reaction (column in the matrix), with negative entries representing reactant consumption and positive entries indicating product formation. Due to mass conservation, the sum of positive entries in each column must sum to 1, and the sum of negative entries must sum to -1. External reactants are denoted by the prefix “x_”. By default, the last row “P” corresponds to the total protein concentration, while the last column corresponds to the ribosome reaction “r” that produces protein.
- Michaelis constant matrix **K**: Contains Michaelis constants in *gL*^*−*1^, organized with rows and columns in the same order as in **M**. Every non-zero entry in **M** must have a corresponding positive entry in **K**, since the reactant is then involved in the reaction.
- Activation constants matrix **KA** and inhibition constants matrix **KI**: expressed in *gL*^*−*1^ and organized with rows and columns in the same order as in **M**. Zero values for individual entries indicate no inhibition nor activation effects for the corresponding internal reactant *ι* on the corresponding reaction *α*.
- Turnover number matrix **k**_cat_: organized in two rows, the first containing the forward-direction turnover numbers (**k**_cat,f_), while the second containing the backward-direction values (**k**_cat,b_), with columns ordered as in matrix **M** and in unit of mass of product produced per mass of catalyst protein per hour (in *h*^*−*1^). An irreversible reaction *α* has the corresponding backward turnover number 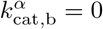.
- Matrix of environment conditions: defines parameters for a growth conditions in each columns. The first row “rho” specifies cell density *ρ* in *gL*^*−*1^. Subsequent rows contain external reactant concentrations **a** in *gL*^*−*1^, with the number of rows corresponding to the number of external reactants (with “x_” prefix) in **M**.

GBA Solver provides a downloadable GBA model template as a reference for users developing their own models, which can then be edited by any spreadsheet editor. Alternatively, users can build models directly within the “Create Model” section of GBA Solver, by filling the corresponding matrices presented each on its own tab. The application employs the “shinyMatrix” R package [1] to generate and display matrices across model tabs, with custom JavaScript modifications that enhance usability.

The model creation process begins by specifying the number of reactants and reactions, followed by entering their corresponding labels. The system automatically designates the label “r” (corresponding to the ribosome reaction) for the final column and the label “P” (corresponding to total protein) for the final row in the matrices **M, K, KI** and **KA**. By default, the first columns in those matrices are reserved to transport reactions, so all columns are ordered as transport, enzymatic, and ribosome reactions. To help with the creation of new models, the tab for each kinetic parameter displays collapsible cards containing relevant data from the BRENDA database [2], with an additional option to query and filter parameters based on organism, EC numbers, and enzyme substrates.

In the “Condition” tab, users input the number of external reactants and “rho” for cell density. After uploading or creating a GBA model, the application presents a comprehensive preview of the input data. This allows users to review their model thoroughly and make direct modifications within the interface. Once finalized, users can export their refined model in ODS format for future use, supporting an iterative development approach that progressively enhances model performance and accuracy.

## III. MODEL VALIDATION AND NUMERICAL OPTIMIZATION

Once a model is loaded, users can validate its compliance with the GBA framework requirements with the “Check Model” button. This comprehensive validation includes:

- Data integrity: verifies that no missing (NA) or non-numeric values exist throughout the model;
- Dimensional consistency: confirms that all matrices maintain consistent dimensions, with matching numbers of reactions and reactants;
- Non-negative value verification: validates that all external concentrations are non-negative and that cell density is positive;
- Michaelis constant validation: identifies instances where substrates (negative entries in the matrix M) have Michaelis constants incorrectly set to zero. If these errors are detected, the system issues a warning and automatically applies the default low value of 0.1 *gL*^*−*1^ to these parameters. Should any other validation criteria fail, the application displays a specific error message directing users to reset their session and revise the problematic aspects of their model.

Upon successful validation, the system notifies users that their model meets all requirements, enabling them to proceed to subsequent analysis steps. This validation workflow ensures that only properly formatted models enter the computational pipeline.

Users initiate the numerical solution of the GBA model by selecting the “Run” button. The optimization uses the nloptr R package [30], which implements the augmented Lagrangian method for non-linear optimization via the auglag function. This method allows users to choose from three specialized local solvers; by default GBA Solver uses SLSQP (Sequential Least Squares Quadratic Programming) which is in general the best suited for smaller models up to 10 reactions, but users can also change to LBFGS (Low-storage BFGS) and MMA (Method of Moving Asymptotes) solvers with the “Advanced option” button. As for any other nonlinear optimization problem, it is in principle not possible to determine beforehand which numerical method will work best for each particular GBA model, and exploration is necessary in each case to find the faster and most accurate local solver. During optimization, a dynamic progress bar tracks completion for each growth condition, providing users with real-time estimates of the total processing time. For reference, the following GBA model example in Fig.1 has 5 reactions and takes at the order of 0.1 second to be optimized per condition.

**FIG. 1.**
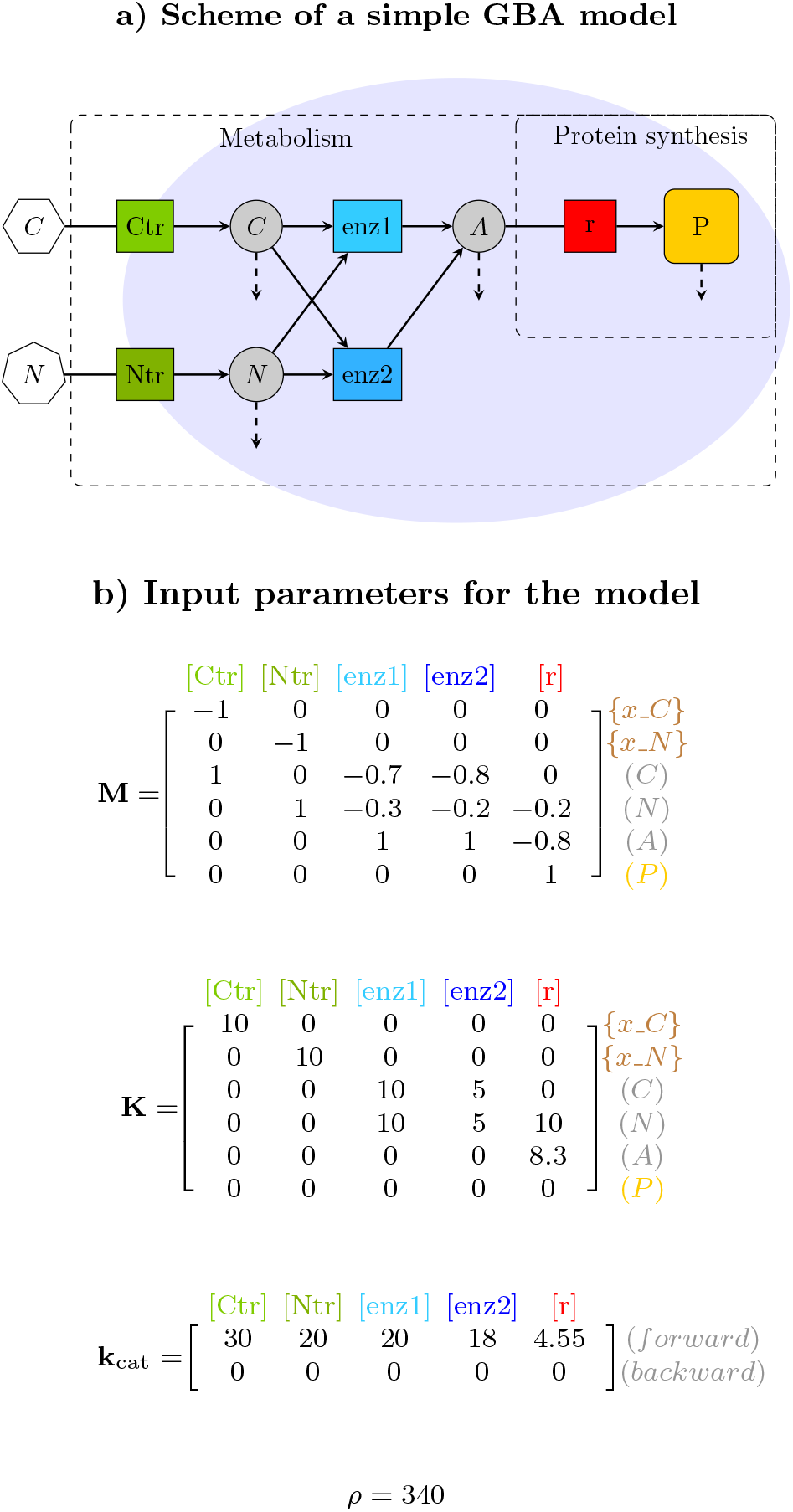
Example of a simple GBA model with 5 irreversible reactions following the Michaelis-Menten kinetics. a) Schematic representation with reactions (squares) and their relationship (solid arrows) with reactants (internal metabolites as circles, total protein “P” as a rounded square, and external reactants as polygons). The GBA framework includes the dilution by growth of each internal reactant (metabolites and total protein) as indicated by dashed arrows, and the necessary protein synthesis by the ribosome reaction “r” to catalyze all reactions. b) Parameters defining the model: 1)mass fraction matrix (**M**) of external and internal reactants, specifying the consumption and production of each reactant (rows) by each reaction (columns), with negative values indicating consumption and positive values indicating production, 2) the matrix of Michaelis constants, and 3) the density *ρ*. Note that since all backward turnover numbers **k**_cat,b_ are zero, as so are the activation and inhibition matrices (not shown), the kinetics of all reactions (2) reduce to a simple Michaelis-Menten rate law.

The numerical solutions across all simulation conditions are presented as interactive visualizations using the apexcharter package [23]. These visualizations include growth rates, protein concentrations, protein fractions, reaction fluxes, and metabolite concentrations. Each interactive plot supports zooming and panning to enable researchers to investigate specific trends. Users can export these visualizations in both SVG and JPEG formats. GBA Solver also integrates d3flux [14] to automatically generate interactive metabolic pathway diagrams directly within the web interface. These pathway visualizations dynamically represent the GBA analysis results from the first growth condition, with line thicknesses and node sizes proportionally reflecting flux magnitudes and concentration values, respectively.

## IV. DISCUSSION

GBA Solver is a user-friendly web application for growth balance analysis (GBA) that increases accessibility and reproducibility in theoretical studies of optimal cell resource allocation at balanced growth. This platform provides an intuitive interface built with R/Shiny, supplemented by HTML, CSS, and JavaScript to enhance the user experience. The application allows users to construct and modify GBA models formatted as spreadsheets (here, .ods files), a familiar open-source format for many researchers. This design choice simplifies the process of inputting and managing model parameters and growth environments without requiring programming skills. GBA Solver also streamlines the incorporation of kinetic parameters into models by integrating data tables retrieved from the BRENDA enzyme database [2].

While the GBA formalism itself imposes no limit on model size, the numerical solution of the corresponding nonlinear optimization problem for large models is still out of reach with current available solvers. For this reason, GBA Solver is at the moment limited to coarsegrained models up to 20 reactions. Cell models up to that size can still capture major resource allocation patterns [5, 10, 13, 20, 26, 29], while resulting in feasible and relatively fast solutions for GBA models with the currently available solvers. This current limitation can in principle be removed by incorporating numerical solvers especially developed for the GBA problem, so larger, more detailed GBA models might be also accepted in this platform.

## VI. ACKNOWLEDGMENTS

We thank Martin J. Lercher for many valuable discussions and suggestions. We are also grateful to Thomas Spitzlei for his support in deploying the application online.

## VII. AUTHOR CONTRIBUTIONS STATEMENT

S.G. and H.D. conceived the GBA Solver platform and wrote the manuscript. S.G. developed and implemented the functionalities and user interface. H.D. supervised the project, provided conceptual guidance, contributed to the user experience improvements.

## IX. DATA AVAILABILITY

GBA Solver is a web server freely accessible without login requirement at https://cellgrowthsim.com/.

The source code is available at https://gitlab.cs.uni-duesseldorf.de/general/ccb/GBASolver.

## References

[1] Andreas Neudecker. shinyMatrix: Shiny Matrix InputArchive Network Pages: 0.8.0.

[2] Antje Chang, Lisa Jeske, Sandra Ulbrich, Julia Hofmann, Julia Koblitz, Ida Schomburg, Meina Neumann-Schaal, Dieter Jahn, and Dietmar Schomburg. BRENDA, the ELIXIR core data resource in 2021: new developments and updates. Nucleic Acids Research, 49(D1):D498– D508, January 2021.

[3] Winston Chang, Joe Cheng, Jj Allaire, Carson Sievert, Barret Schloerke, Yihui Xie, Jeff Allen, Jonathan McPherson, Alan Dipert, and Barbara Borges. shiny: Web Application Framework for R, December 2012. Institution: Comprehensive R Archive Network Pages: 1.9.1.

[4] Kiri Choi, J. Kyle Medley, Matthias König, Kaylene Stocking, Lucian Smith, Stanley Gu, and Herbert M. Sauro. Tellurium: An extensible python-based modeling environment for systems and synthetic biology. Biosystems, 171:74–79, September 2018.

[5] Dieu Thi Doan, Manh Dat Hoang, Anna-Lena Heins, and Andreas Kremling. Applications of Coarse-Grained Models in Metabolic Engineering. Frontiers in Molecular Biosciences, 9:806213, March 2022.

[6] Hugo Dourado and Martin J. Lercher. An analytical theory of balanced cellular growth. Nature Communications, 11(1):1226, March 2020.

[7] Hugo Dourado, Wolfram Liebermeister, Oliver Ebenhöh, and Martin J. Lercher. Mathematical properties of optimal fluxes in cellular reaction networks at balanced growth. PLOS Computational Biology, 19(6):e1011156, June 2023. Number: 6.

[8] David W. Erickson, Severin J. Schink, Vadim Patsalo, James R. Williamson, Ulrich Gerland, and Terence Hwa. A global resource allocation strategy governs growth transition kinetics of Escherichia coli. Nature, 551(7678):119–123, November 2017. Publisher: Nature Publishing Group.

[9] Fabian Fröhlich, Daniel Weindl, Yannik Schälte, Dilan Pathirana, Lukasz Paszkowski, Glenn Terje Lines, Paul Stapor, and Jan Hasenauer. AMICI: high-performance sensitivity analysis for large ordinary differential equation models. Bioinformatics, 37(20):3676–3677, October 2021.

[10] Sajjad Ghaffarinasab, Martin J. Lercher, and Hugo Dourado. Growth balance analysis models of cyanobacteria for understanding resource allocation strategies, April 2023.

[11] Archana Hari and Daniel Lobo. Fluxer: a web application to compute, analyze and visualize genomescale metabolic flux networks. Nucleic Acids Research, 48(W1):W427–W435, July 2020.

[12] Stefan Hoops, Sven Sahle, Ralph Gauges, Christine Lee, Juürgen Pahle, Natalia Simus, Mudita Singhal, Liang Xu, Pedro Mendes, and Ursula Kummer. COPASI—a COmplex PAthway SImulator. Bioinformatics, 22(24):3067– 3074, December 2006.

[13] Sheng Hui, Josh M Silverman, Stephen S Chen, David W Erickson, Markus Basan, Jilong Wang, Terence Hwa, and James R Williamson. Quantitative proteomic analysis reveals a simple strategy of global resource allocation in bacteria. Molecular Systems Biology, 11(2):784, February 2015.

[14] Peter St. John. D3flux: D3.js-based visualizations of cobrapy metabolic models. GitHub repository, 2021. Accessed: December 14, 2024.

[15] Zachary A. King, Andreas Dräger, Ali Ebrahim, Nikolaus Sonnenschein, Nathan E. Lewis, and Bernhard O. Palsson. Escher: A Web Application for Building, Sharing, and Embedding Data-Rich Visualizations of Biological Pathways. PLoS Computational Biology, 11(8):1–13, 2015.

[16] Wolfram Liebermeister and Edda Klipp. Bringing metabolic networks to life: convenience rate law and thermodynamic constraints. Theoretical Biology and Medical Modelling, 3(1):41, December 2006.

[17] Greg Lin. reactable: Interactive Data Tables for R, November 2019. Institution: Comprehensive R Archive Network Pages: 0.4.4.

[18] Zhitao Mao, Qianqian Yuan, Haoran Li, Yue Zhang, Yuanyuan Huang, Chunhe Yang, Ruoyu Wang, Yongfu Yang, Yalun Wu, Shihui Yang, Xiaoping Liao, and Hongwu Ma. CAVE: a cloud-based platform for analysis and visualization of metabolic pathways. Nucleic Acids Research, 51(W1):W70–W77, July 2023.

[19] Nikolay Martyushenko and Eivind Almaas. Model-Explorer - software for visual inspection and inconsistency correction of genome-scale metabolic reconstructions. BMC Bioinformatics, 20(1):56, December 2019.

[20] Douwe Molenaar, Rogier Van Berlo, Dick De Ridder, and Bas Teusink. Shifts in growth strategies reflect tradeoffs in cellular economics. Molecular Systems Biology, 5(323):1–10, 2009. Publisher: Nature Publishing Group.

[21] Brett G. Olivier and Jacky L. Snoep. Web-based kinetic modelling using JWS Online. Bioinformatics, 20(13):2143–2144, September 2004.

[22] Jeffrey D. Orth, Ines Thiele, and Bernhard Ø Palsson. What is flux balance analysis? Nature Biotechnology, 28(3):245–248, March 2010. Publisher: Nature Publishing Group.

[23] Victor Perrier and Fanny Meyer. apexcharter: Create Interactive Chart with the JavaScript ‘ApexCharts’ Library, July 2019. Institution: Comprehensive R Archive Network Pages: 0.4.4.

[24] R Core Team. R: A Language and Environment for Statistical Computing. R Foundation for Statistical Computing, Vienna, Austria, 2024.

[25] J Schaff, CC Fink, B Slepchenko, JH Carson, and LM Loew. A general computational framework for modeling cellular structure and function. Biophysical Journal, 73(3):1135–1146, September 1997.

[26] Matthew Scott, Carl W. Gunderson, Eduard M. Mateescu, Zhongge Zhang, and Terence Hwa. Interdependence of Cell Growth and Gene Expression: Origins and Consequences. Science, 330(6007):1099–1102, November 2010.

[27] Bilal Shaikh, Gnaneswara Marupilla, Mike Wilson, Michael L Blinov, Ion I Moraru, and Jonathan R Karr. RunBioSimulations: an extensible web application that simulates a wide range of computational modeling frameworks, algorithms, and formats. Nucleic Acids Research, 49(W1):W597–W602, July 2021.

[28] Sven Thiele, Axel Von Kamp, Pavlos Stephanos Bekiaris, Philipp Schneider, and Steffen Klamt. CNApy: a Cell-NetAnalyzer GUI in Python for analyzing and designing metabolic networks. Bioinformatics, 38(5):1467–1469, February 2022. Number: 5.

[29] Andrea Y. Weiße, Diego A. Oyarzún, Vincent Danos, and Peter S. Swain. Mechanistic links between cellular trade-offs, gene expression, and growth. Proceedings of the National Academy of Sciences, 112(9), March 2015.

[30] Jelmer Ypma and Steven G. Johnson. nloptr: R Interface to NLopt, June 2011. Institution: Comprehensive R Archive Network Pages: 2.1.1.

